# dyngen: a multi-modal simulator for spearheading new single-cell omics analyses

**DOI:** 10.1101/2020.02.06.936971

**Authors:** Robrecht Cannoodt, Wouter Saelens, Louise Deconinck, Yvan Saeys

**Affiliations:** Ghent University

## Abstract

We present dyngen, a novel, multi-modal simulation engine for studying dynamic cellular processes at single-cell resolution. dyngen is more flexible than current single-cell simulation engines, and allows better method development and benchmarking, thereby stimulating development and testing of novel computational methods. We demonstrate its potential for spearheading novel computational methods on three novel applications: aligning cell developmental trajectories, single-cell regulatory network inference and estimation of RNA velocity.

## Main text

Single-cell simulation engines are becoming increasingly important for testing and benchmarking computational methods, a pressing need in the widely expanding field of single-cell biology. Complementary to real biological data, synthetic data provides a valuable alternative where the actual ground truth is completely known and thus can be compared to, in order to make quantitative evaluations of computational methods that aim to reconstruct this ground truth [1]. In addition, simulation engines are more flexible when it comes to stress-testing computational methods, for example by varying the parameters of the simulation, such as the amount of noise, samples, and cells measured, allowing benchmarking of methods over a wide range of possible scenarios. In this way, they can even guide the design of real biological experiments, finding out the best conditions to be used as input for subsequent computational pipelines. Another, more experimental use of simulation engines is their important role in spearheading the development of novel computational methods, possibly even before real data is available. In this way, simulation engines can be used to assess the value of novel experimental protocols or treatments. Simulation engines are also increasingly important when it comes to finding alternatives to animal models, for example for drug testing and precision medicine. In such scenarios, cellular simulations can act as digital twins, offering unlimited experimentation *in silico* [2].

Here, we introduce dyngen, a novel multi-modal simulator of dynamic biological processes at single-cell resolution (Figure 1). dyngen uses Gillespie’s stochastic simulation algorithm [3] to simulate gene regulation, splicing, and translation at a single-molecule level. Other generators of scRNA-seq data (e.g. splatter [1], powsimR [4], PROSSTT [5] and SymSim [6]) have already been used extensively to explore the strengths and weaknesses of computational tools, both by method developers [7, 8, 9, 10] and independent benchmarkers [11, 12, 13]. However, a limitation of these existing simulators is that they would require significant methodological alterations to add additional modalities or experimental conditions (Table S1).

**Figure 1:**
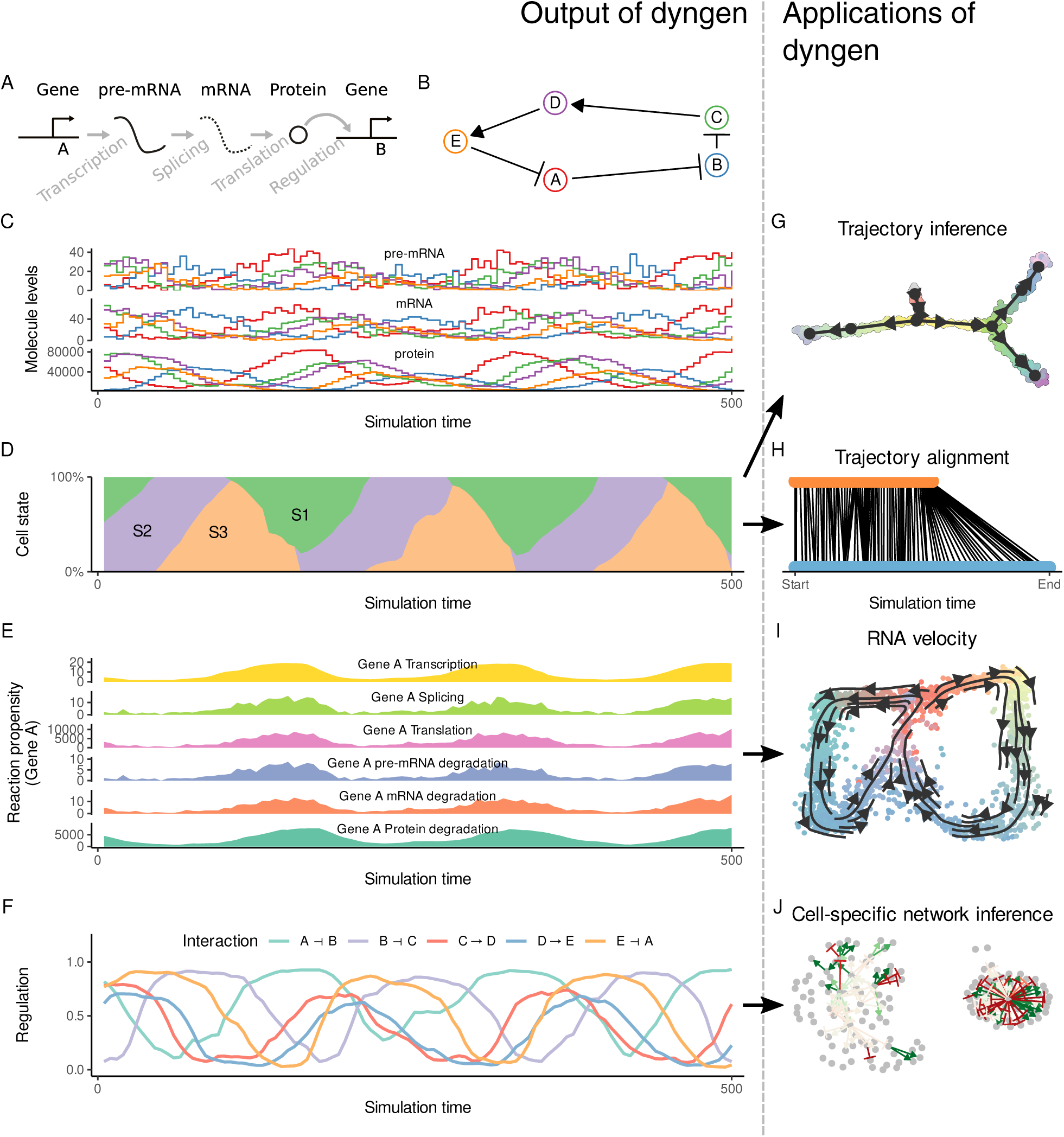
Showcase of dyngen functionality. **A:** Changes in abundance levels are driven strictly by gene regulatory reactions. **B:** The input Gene Regulatory Network (GRN) is defined such that it models a dynamic process of interest. **C:** The reactions define how abundance levels of molecules change at any particular time point. **D:** Firing many reactions can significantly alter the cellular state over time. **E:** dyngen keeps track of the likelihood of a reaction firing during small intervals of time, called the propensity, as well as the actual number of firings. **F:** Similarly, dyngen can also keep track of the regulatory activity of every interaction. **G:** A benchmark of trajectory inference methods has already been performed using the cell state ground-truth. **H:** The cell state ground-truth enables evaluating trajectory alignment methods [12]. **I:** The reaction propensity ground-truth enables evaluating RNA velocity methods. **J:** The cellwise regulatory network ground-truth enables evaluating cell-specific gene regulatory network inference methods.

dyngen was designed to include all of these functionalities and more by design. Its methodology allows tracking many layers of information throughout the simulation, including the abundance of any molecule in the cell, the progression of the cell along a dynamic process, and the activation strength of individual regulatory interactions. dyngen can simulate a large variety of dynamic processes (e.g. cyclic, branching, disconnected) as well as a broad range of experimental conditions (e.g. batch effects and time-series, perturbation and knockdown experiments). The fine-grained controls over simulation parameters allow dyngen to be applicable to a broad range of use-cases to simulate dynamic biological processes. The original design of dyngen was motivated by the plethora of methods available for trajectory inference, where dyngen allowed the first large-scale benchmarking of such methods [12, 14].

Here, we highlight the novel functionality of dyngen by evaluating three novel types of computational approaches for which no simulation engines exist yet: single-cell network inference, trajectory alignment and RNA velocity (Figure 2). We emphasize that our main aim here is to illustrate the potential of dyngen for these evaluations, rather than performing large-scale benchmarking, which would require assessing many more quantitative and qualitative aspects of each method [15].

**Figure 2:**
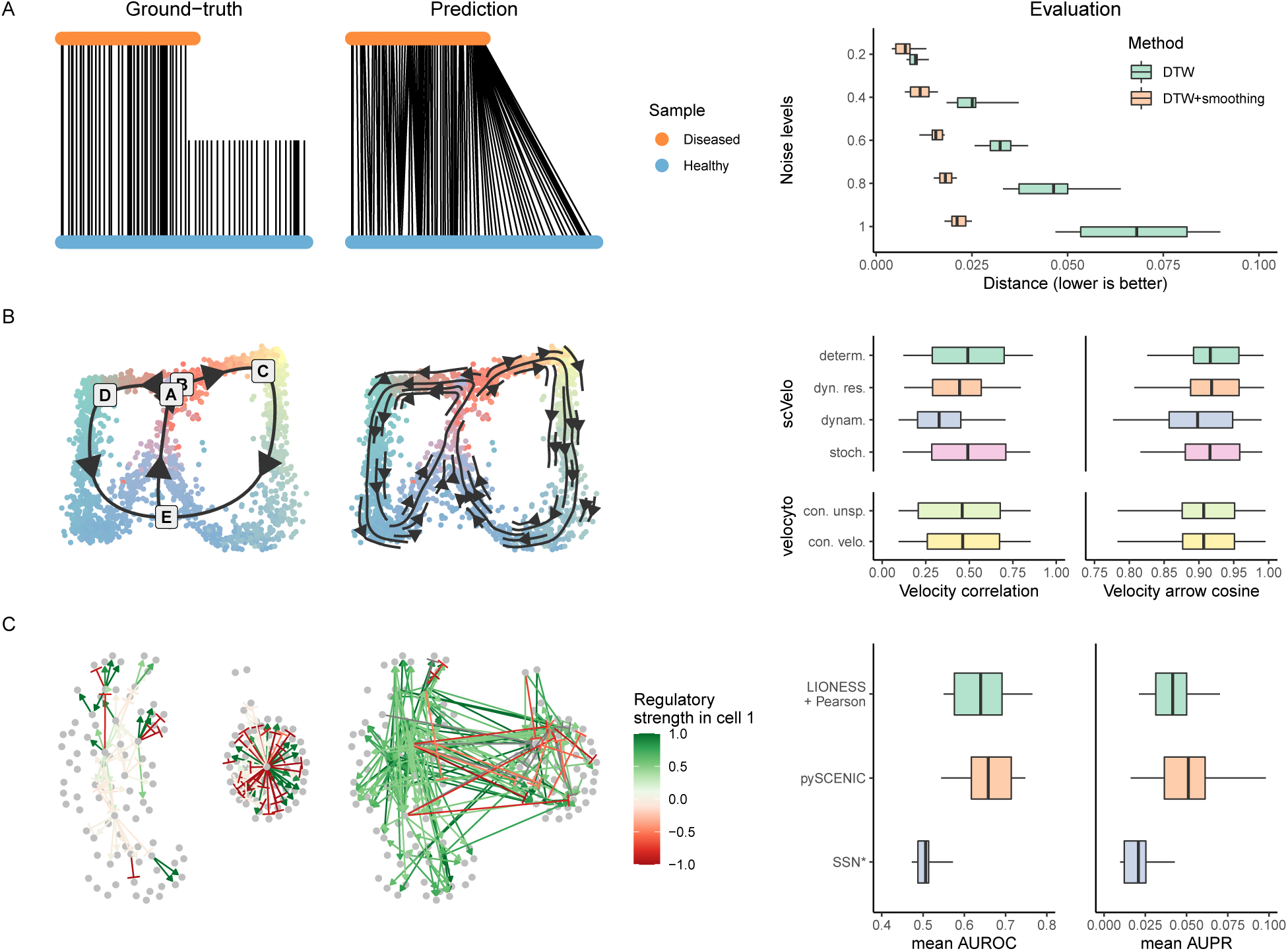
dyngen provides ground-truth data for a variety of applications (left), which can be used to quantitatively evaluate methods (right). **A:** Trajectory alignment aligns two trajectories between samples. dyngen can simulate different scenarios in which alignment is necessary, such as a premature stop as shown here. We compared two versions of dynamic time warping (DTW): normal DTW aligns all individual cells, while DTW+smoothing first smooths the expression data and then uses DTW to align the smoothed cells. **B:** RNA velocity calculates for each cell the direction in which the expression of each gene is moving. We evaluated scVelo and velocyto by comparing these vectors with the known velocity vector (velocity correlation) and with the known direction of the cellular trajectory in a dimensionality reduction (velocity arrow cosine). **C:** Cell-specific network inference (CSNI) predicts the regulatory network of every individual cell. We evaluate each cell-specific regulatory network with typical metrics for network inference: the Area Under the Receiver Operating Characteristics-curve (AUROC) and Area Under the Precision Recall-curve (AUPR). We evaluate three CSNI methods by computing the mean AUROC and AUPR across all cells.

### Trajectory alignment

methods align trajectories from different samples and allow studying the differences between the different trajectories. For example, by comparing the transcriptomic profiles of cells from a diseased patient to a healthy control, it might be possible to detect transcriptomics differences (differential expression) of particular cells along a developmental process, or to detect an early stop of the trajectory of the diseased patient. Currently, trajectory alignment is limited to aligning linear trajectories, though other topologies of trajectory could be aligned as well. Dynamic Time Warping (DTW) [16] is a method designed for aligning temporal sequences for speech recognition but has since been used to compare gene expression kinetics from many different biological processes [17, 18, 19, 20]. We evaluate the performance of DTW by simulating two linear trajectories with slightly different simulation kinetics and assessing the accuracy of the alignment by DTW. We compare the performance of DTW against a version of DTW where gene expression is first smoothed, for varying degrees of noise added to the gene expression (Figure S1). We observe that DTW+smoother performs significantly better than the unsmoothed version across all levels of noise.

### RNA velocity

methods use the relative ratio between pre-mRNA and mature mRNA reads to predict the velocity at which the RNA expression of genes is increasing or decreasing [21, 22]. Already two algorithms are currently available for estimating the RNA velocity vector from spliced and unspliced counts: velocyto [22] and scvelo [23]. Yet, to date, no quantitative assessment of their accuracy has been performed, mainly due to the difficulty in obtaining real ground-truth data to do so. In contrast, the ground-truth RNA velocity can be easily extracted from a dyngen, as it is possible to store the rate at which mRNA molecules are being transcribed and degraded (called the propensity) at any particular point in time. We executed scvelo and velocyto (with 6 different parameter settings in total) on 102 datasets with varying degrees of difficulty (easy, medium, hard) and a variety of backbones (including linear, bifurcating, cyclic, disconnected). We evaluated the predictions using two metrics (Figure S2), one which directly compares the predicted RNA velocity of each gene with the ground-truth RNA velocity (called the “velocity correlation”), and one which compares the direction of the ground-truth trajectory embedded in a dimensionality reduction with the average RNA velocity of cells in that neighbourhood (called the “velocity arrow cosine”). We found that in almost all cases, both velocyto and scvelo obtained high velocity arrow cosine, meaning that the overview obtained by embedding RNA velocity arrows in a 2D or 3D dimensionality reduction allows users to correctly identify the general progression of cells. However, depending on the difficulty of the dataset, the correlation between the predicted RNA velocity and the ground-truth RNA velocity varies between high (>0.75) to low (<0.25). For this particular metric, the dynamic estimation of velocyto performs significantly worse than any of the other prediction methods.

### Cell-specific network inference

(CSNI) methods predict not only which transcription factors regulate which target genes, but also aim to identify how active each interaction is in each of the cells, since interactions can be turned off and on depending on the cellular state. While a few pioneering CSNI approaches have already been developed [24, 25, 26], a quantitative assessment of the performance is until now lacking. This is not surprising, as neither real nor in silico datasets of cell-specific or even cell-type-specific interactions exist that are large enough so that it can be used as a ground-truth for evaluating CSNI methods. Extracting the ground-truth dynamic network in dyngen is straightforward though, given that we can calculate how target gene expression would change without the regulator being present. We used this ground-truth to compare the performance of three CSNI methods (Figure S3): LIONESS [25], SSN [26] and SCENIC [24]. We calculated the AUROC and AUPR score for each cell individually. Computing the mean AUROC and AUPR per dataset showed that pySCENIC significantly outperforms LIONESS + Pearson, which in turn outperforms SSN*.

In summary, dyngen’s single-cell simulations can be used to evaluate common single-cell omics computational methods such as clustering, batch correction, trajectory inference, and network inference. However, the framework is flexible enough to be adaptable to a broad range of applications, including methods that integrate clustering, network inference, and trajectory inference. In this respect, dyngen may promote the development of new tools in the single-cell field similarly as other simulators have done in the past [27, 28]. Additionally, one could anticipate technological developments in single-cell multi-omics. In this way, dyngen allows designing and evaluating the performance and robustness of new types of computational analyses before experimental data becomes available, comparing which experimental protocol is the most cost-effective in producing qualitative and robust results in downstream analysis. One major assumption of dyngen is that cells are regarded as standalone entities that are well mixed. Splitting up the simulation space into separate subvolumes could pave the way to better study key cellular processes such as cell division, intercellular communication, and migration [29].

## Availability

dyngen is available as an open-source R package on CRAN at cran.r-project.org/package=dyngen. The analyses performed in this manuscript are available on GitHub at github.com/dynverse/dyngen_manuscript.

## Author contributions

- W.S. and R.C. designed the study.
- R.C., W.S., and L.D. performed the experiments and analysed the data.
- R.C. and W.S. implemented the dyngen software package.
- R.C., W.S., L.D., and Y.S. wrote the manuscript.
- Y.S. supervised the project.

## Methods

The workflow to generate *in silico* single-cell data consists of six main steps (Figure S4).

### Defining the module network

One of the main processes involved in cellular dynamic processes is gene regulation, where regulatory cascades and feedback loops lead to progressive changes in expression and decision making. The exact way a cell chooses a certain path during its differentiation is still an active research field, although certain models have already emerged and been tested *in vivo*. One driver of bifurcation is mutual antagonism, where two genes strongly repress each other [30, 31], forcing one of the two to become inactive [32]. Such mutual antagonism can be modelled and simulated [33, 34]. Although the two-gene model is simple and elegant, the reality is frequently more complex, with multiple genes (grouped into modules) repressing each other [35].

To start a dyngen simulation, the user needs to define a module network. The module network describes how sets of genes regulate each other and is what mainly determines which dynamic processes occur within the simulated cells.

A module network consists of modules connected together by regulatory interactions, which can be either up- or down-regulating. A module may have basal expression, which means genes in this module will be transcribed without the presence of transcription factor molecules. A module marked as “active during the burn phase” means that this module will be allowed to generate expression of its genes during an initial warm-up phase (See section). At the end of the dyngen process, cells will not be sampled from the burn phase simulations. Interactions between modules have a strength (which is a positive integer) and an effect (+1 for upregulating, -1 for downregulating).

Several examples of module networks are given in Figure S5. A simple chain of modules (where one module upregulates the next) results in a *linear* process. By having the last module repress the first module, the process becomes *cyclic*. Two modules repressing each other is the basis of a *bifurcating* process, though several chains of modules have to be attached in order to achieve progression before and after the bifurcation process. Finally, a *converging* process has a bifurcation occurring during the burn phase, after which any differences in module regulation is removed.

Note that these examples represent the bare minimum in terms of the number of modules used. Using longer chains of modules is typically desired. In addition, the fate decisions made in this example of a bifurcation is reversible, meaning cells can be reprogrammed to go down a different differentiation path. If this effect is undesirable, more safeguards need to be put in place to prevent reprogramming from occurring.

### Generating the gene regulatory network

The GRN is generated based on the given module network in four main steps (Figure S6).

**Step 1, sampling the transcription factors (TF)**. The TFs are the main drivers of the molecular changes in the simulation. The user provides a backbone and the number of TFs to generate. Each TF is assigned to a module such that each module has at least *x* parameters (default *x* = 1). A TF inherits the ‘burn’ and ‘basal expression’ from the module it belongs to.

**Step 2, generating the TF interactions**. Let each TF be regulated according to the interactions in the backbone. These interactions inherit the effect, strength, and independence parameters from the interactions in the backbone. A TF can only be regulated by other TFs or itself.

**Step 3, sampling the target subnetwork**. A user-defined number of target genes are added to the GRN. Target genes are regulated by a TF or another target gene, but are always downstream of at least one TF. To sample the interactions between target genes, one of the many FANTOM5 [36] GRNs is sampled. The currently existing TFs are mapped to regulators in the FANTOM5 GRN. The targets are drawn from the FANTOM5 GRN weighted by their page rank value, to create an induced GRN. For each target, at most *x* regulators are sampled from the induced FANTOM5 GRN (default *x* = 5). The interactions connecting a target gene and its regulators are added to the GRN.

**Step 4, sampling the housekeeping subnetwork**. Housekeeping genes are completely separate from any TFs or target genes. A user-defined set of housekeeping genes is also sampled from the FANTOM5 GRN. The interactions of the FANTOM5 GRN are first subsampled such that the maximum in-degree of each gene is *x* (default *x* = 5). A random gene is sampled and a breadth-first-search is performed to sample the desired number of housekeeping genes.

### Convert gene regulatory network to a set of reactions

Simulating a cell’s GRN makes use of a stochastic framework which tracks the abundance levels of molecules over time in a discrete quantity. For every gene *G*, the abundance levels of three molecules are tracked, namely of corresponding pre-mRNAs, mature mRNAs and proteins, which are represented by the terms x_*G*_, y*G* and z_*G*_ respectively. The GRN defines how a reaction affects the abundance levels of molecules and how likely it will occur. Gibson and Bruck [37] provide a good introduction to modelling gene regulation with stochastic frameworks, on which many of the concepts below are based.

For every gene in the GRN a set of reactions are defined, namely transcription, splicing, translation, and degradation. Each reaction consists of a propensity function – a formula *f*(.) to calculate the probability *f*(.) × d*t* of it occurring during a time interval d*t* – and the effect – how it will affect the current state if triggered.

The effects of each reaction mimic the respective biological processes (Table S2, middle). Transcription of gene *G* results in the creation of a single pre-mRNA molecule x_*G*_. Splicing turns one pre-mRNA x_*G*_ into a mature mRNA x_*G*_. Translation uses a mature mRNA y*G* to produce a protein z_*G*_. Pre-mRNA, mRNA and protein degradation results in the removal of a x_*G*_, y*G*, and z_*G*_ molecule, respectively.

The propensity of all reactions except transcription are all linear functions (Table S2, right) of the abundance level of some molecule multiplied by a per-gene constant (Table S3). The propensity of transcription of a gene *G* depends on the abundance levels of its TFs. The per-gene and per-interaction constants are based on the median reported production-rates and half-lives of molecules measured of 5000 mammalian genes [38], except that the transcription rate has been amplified by a factor of 10.

The propensity of the transcription of a gene *G* is inspired by thermodynamic models of gene regulation [39], in which the promoter of *G* can be bound or unbound by a set of *N* transcription factors *H*_*i*_. Let *f*(z_1_, z_2_,…, z_*N*_) denote the propensity function of *G*, in function of the abundance levels of the transcription factors. The following subsections explain and define the propensity function when *N* = 1, *N* = 2, and finally for an arbitrary *N*.

### Propensity of transcription when *N* = 1

In the simplest case when *N* = 1, the promoter can be in one of two states. In state *S*_0_, the promoter is not bound by any transcription factors, and in state *S*_1_ the promoter is bound by *H*_1_. Each state *S*_*j*_ is linked with a relative activation *α*_*j*_, a number between 0 and 1 representing the activity of the promoter at this particular state. The propensity function is thus equal to the expected value of the activity of the promoter multiplied by the pre-mRNA production rate of *G*.

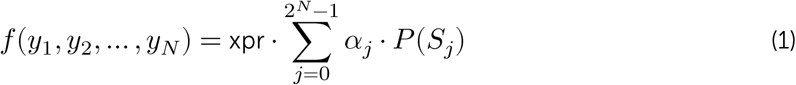

For *N* = 1, *P* (*S*_1_) is equal to the Hill equation, where *k*_*i*_ represents the concentration of *H*_*i*_ at half-occupation and *n*_*i*_ represents the Hill coefficient. Typically, *n*_*i*_ is between [1,10]

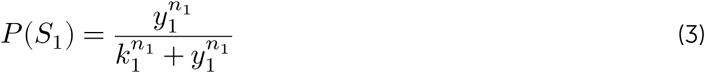

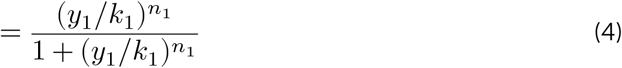

The Hill equation can be simplified by letting 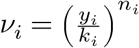.

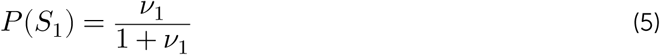

Since *P* (*S*_0_) = 1 − *P* (*S*_1_), the activation function is formulated and simplified as follows.

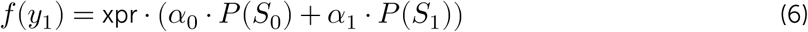

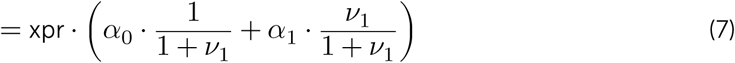

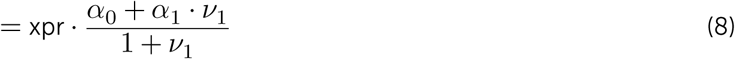

### Propensity of transcription when *N* = 2

When *N* = 2, there are four states *S*_*j*_. The relative activations *α*_*j*_ can be defined such that *H*_1_ and *H*_2_ are independent (additive) or synergistic (multiplicative). In order to define the propensity of transcription *f*(.), the Hill equation *P* (*S*_*j*_) is extended for two transcription factors.

Let *w*_*j*_ be the numerator of *P* (*S*_*j*_), defined as the product of all transcription factors bound in that state:

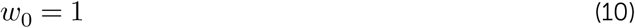

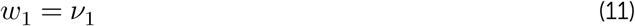

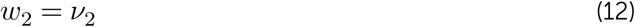

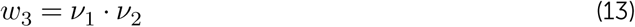

The denominator of *P* (*S*_*j*_) is then equal to the sum of all *w*_*j*_. The probability of state *S*_*j*_ is thus defined as:

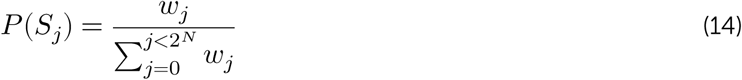

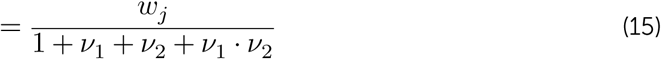

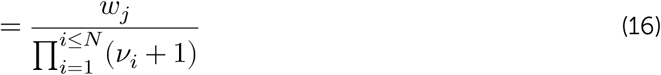

Substituting *P* (*S*_*j*_) and *w*_*j*_ into *f*(.) results in the following equation:

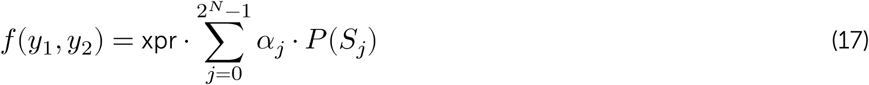

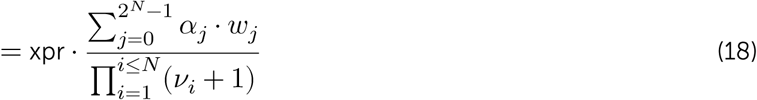

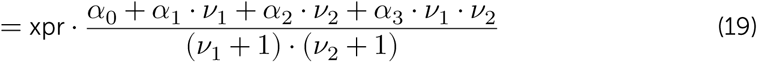

### Propensity of transcription for an arbitrary *N*

For an arbitrary *N*, there are 2^*N*^ states *S*_*j*_. The relative activations *α*_*j*_ can be defined such that *H*_1_ and *H*_2_ are independent (additive) or synergistic (multiplicative). In order to define the propensity of transcription *f*(.), the Hill equation *P* (*S*_*j*_) is extended for *N* transcription factors.

Let *w*_*j*_ be the numerator of *P* (*S*_*j*_), defined as the product of all transcription factors bound in that state:

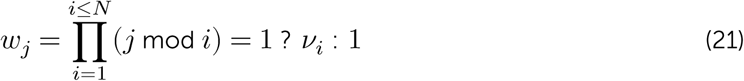

The denominator of *P* (*S*_*j*_) is then equal to the sum of all *w*_*j*_. The probability of state *S*_*j*_ is thus defined as:

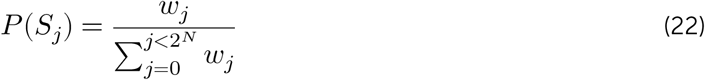

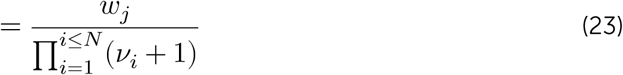

Substituting *P* (*S*_*j*_) into *f*(.) yields:

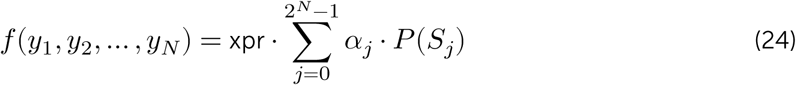

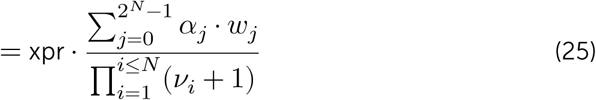

### Propensity of transcription for a large *N*

For large values of *N*, computing *f*(.) is practically infeasible as it requires performing 2^*N*^ summations. In order to greatly simplify *f*(.), *α*_*j*_ could be defined as 0 when one of the regulators inhibits transcription and 1 otherwise.

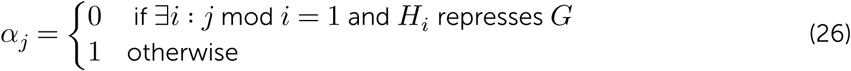

Substituting equation 26 into equation 25 and defining *R* = {1, 2, …, *N*} and *R*^+^ = {*i*|*H*_*i*_ activates *G*} yields the simplified propensity function:

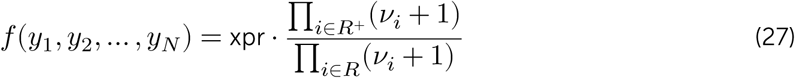

### Independence, synergism and basal expression

The definition of *α*_*j*_ as in equation 26 presents two main limitations. Firstly, since *α*_0_ = 1, it is impossible to tweak the propensity of transcription when no transcription factors are bound. Secondly, it is not possible to tweak the independence and synergism of multiple regulators.

Let ba ∈ [0, 1] denote the basal expression strength *G* (i.e. how much will *G* be expressed when no transcription factors are bound), and sy ∈ [0, 1] denote the synergism of regulators *H*_*i*_ of *G*, the transcription propensity becomes:

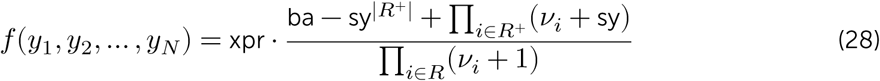

### Simulate single cells

dyngen uses Gillespie’s stochastic simulation algorithm (SSA) [3] to simulate dynamic processes. An SSA simulation is an iterative process where at each iteration one reaction is triggered.

Each reaction consists of its propensity – a formula to calculate the probability of the reaction occurring during an infinitesimal time interval – and the effect – how it will affect the current state if triggered. Each time a reaction is triggered, the simulation time is incremented by 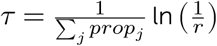 with *r* ∈ *U*(0, 1) and *prop*_*j*_ the propensity value of the *j*th reaction for the current state of the simulation.

GillespieSSA2 is an optimised library for performing SSA simulations. The propensity functions are compiled to C++ and SSA approximations can be used which allow triggering many reactions simultaneously at each iteration. The framework also allows storing the abundance levels of molecules only after a specific interval has passed since the previous census. By setting the census interval to 0, the whole simulation’s trajectory is retained but many of these time points will contain very similar information. In addition to the abundance levels, also the propensity values and the number of firings of each of the reactions at each of the time steps can be retained, as well as specific sub-calculations of the propensity values, such as the regulator activity level *reg*_*G,H*_.

### Simulate experiment

From the SSA simulation we obtain the abundance levels of all the molecules at every state. We need to replicate technical effects introduced by experimental protocols in order to obtain data that is similar to real data. For this, the cells are sampled from the simulations and molecules are sampled for each of the cells. Gene capture rates and library sizes are empirically derived from real datasets as to match real technical variation.

### Sample cells

In this step, *N* cells are sampled the simulations. Two approaches are implemented: sampling from an unsynchronised population of single cells (snapshot) or sampling at multiple time points in a synchronised population (time series).

#### Snapshot

The backbone consists of several states linked together by transition edges with length *L*_*i*_, to which the different states in the different simulations have been mapped (Figure S7A). From each transition, 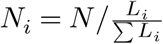 cells are sampled uniformly, rounded such that Σ*N*_*i*_ = *N*.

#### Time series

Assuming that the final time of the simulations is *T*, the interval [0, *T*] is divided into *k* equal intervals of width *w* separated by *k* − 1 gaps of width *g. N*_*i*_ = *N*/*k* cells are sampled uniformly from each interval (Figure S7B), rounded such that Σ*N*_*i*_ = *N*. By default, *k* = 8 and *g* = 0.75. For usual dyngen simulations, 10 ≤ *T* ≤ 20. For larger values of *T, k* and *g* should be increased accordingly.

### Sample molecules

Molecules are sampled from the simulation to replicate how molecules are experimentally sampled. A real dataset is downloaded from a repository of single-cell RNA-seq datasets [40]. For each *in silico* cell *i*, draw its library size *lS*_*i*_ from the distribution of transcript counts per cell in the real dataset. The capture rate *cr*_*j*_ of each *in silico* molecule type *j* is drawn from *N*(1, 0.05). Finally, for each cell *i*, draw *lS*_*i*_ molecules from the multinomial distribution with probabilities *cr*_*j*_ × *ab*_*i,j*_ with *ab*_*i,j*_ the molecule abundance level of molecule *j* in cell *i*.

### Simulating batch effects

Simulating batch effects can be performed in multiple ways. One such way is to perform the first two steps of the creation of a dyngen model (defining the module network and generating the GRN). For each desired batch, create a separate model for which random kinetics are generated and perform all subsequent dyngen steps (convert to reactions, simulate gold standard, simulate single cells, simulate experiment). Since each separate model has different underlying kinetics, the combined output will resemble having batch effects.

### Determining the ground-truth trajectory

To construct the ground-truth trajectory, the user needs to provide the ground-truth state network alongside the initial module network (Figure∼S8). Each edge in the state network specifies which modules are allowed to change in expression in transitioning from one state to another. For each edge, a simulation is run using the end state of an upstream branch as the initial expression vector, and only allowing the modules as predefined by the attribute to change.

As an example, consider the cyclic trajectory shown in Figure∼S8. State S0 begins with an expression vector of all zero values. To simulate the transition from S0 to S1, regulation of the genes in modules A, B and C are turned on. After a predefined period of time, the end state of this transition is considered the expression vector of state S1. To simulate the transition from S1 to S2, regulation of the genes in modules D and E are turned on, while the regulation of genes in module C is turned off. During this simulation, the expression of genes in modules A, B, D, and E is thus allowed to change. The end state of the simulation is considered the expression vector of state S2.

For each of the branches in the state network, an expression matrix and the corresponding progression time along that branch are retained. To map a simulated cell to the ground-truth, the correlation between its expression values and the expression matrix of the ground-truth trajectory is calculated, and the cell is mapped to the position in the ground-truth trajectory that has the highest correlation.

### Determining the cell-specific ground-truth regulatory network

Calculating the regulatory effect of a regulator *R* on a target *T* (Figure S4F) requires determining the contribution of *R* in the propensity function of the transcription of *T* (section) with respect to other regulators. This information is useful, amongst others, for benchmarking cell-specific network inference methods.

The regulatory effect of *R* on *T* at a particular state *S* is defined as the change in the propensity of transcription when *R* is set to zero, scaled by the inverse of the pre-mRNA production rate of *T*. More formally:

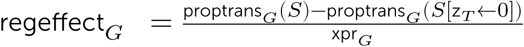

Determining the regulatory effect for all interactions and cells in the dataset yields the complete cell-specific ground-truth GRN. The regulatory effect lies between [−1, 1], where -1 represents complete inhibition of *T* by *R*, 1 represents maximal activation of *T* by *R*, and 0 represents inactivity of the regulatory interaction between *R* and *T*.

### Comparison of cell-specific network inference methods

14 datasets were generated using the 14 different predefined backbones. For every cell in the dataset, the transcriptomics profile and the corresponding cell-specific ground-truth regulatory network was determined (Section).

Several cell-specific NI methods were considered for comparison: SCENIC [24], LIONESS [41, 25], and SSN [26].

LIONESS [25] runs a NI method multiple times to construct cell-specific GRNs. LIONESS first infers a GRN with all of the samples. A second GRN is inferred with all samples except one particular profile. The cell-specific GRN for that particular profile is defined as the difference between the two GRN matrices. This process is repeated for all profiles, resulting in a cell-specific GRN. By default, LIONESS uses PANDA [42] to infer GRNs, but since dyngen does not produce motif data and motif data is required by PANDA, PANDA is inapplicable in this context. Instead, we used the lionessR [43] implementation of LIONESS, which uses by default the Pearson correlation as a NI method. We marked results from this implementation as “LIONESS + Pearson”.

SSN [26] follows, in essence, the exact same methodology as LIONESS except that it specifically only uses the Pearson correlation. It is worth noting that the LIONESS preprint was released before the publication of SSN. Since no implementation was provided by the authors, we implemented SSN in R using basic R and tidyverse functions [44] and marked results from this implementation as “SSN*”.

SCENIC [24] is a pipeline that consists of four main steps. Step 1: classical network inference is performed with arboreto, which is similar to GENIE3 [45]. Step 2: select the top 10 regulators per target. Interactions are grouped together in ‘modules’; each module contains one regulator and all of its targets. Step 3: filter the modules using motif analysis. Step 4: for each cell, determine an activity score of each module using AUCell. As a post-processing of this output, all modules and the corresponding activity scores are combined back into a cell-specific GRN consisting of (cell, regulator, target, score) pairs. For this analysis, the Python implementation of SCENIC was used, namely pySCENIC. Since dyngen does not generate motif data, step 3 in this analysis is skipped.

Cell-specific network inference (CSNI) predicts the regulatory network of every individual cell. We evaluate each cell-specific regulatory network with typical metrics for network inference: the Area Under the Receiver Operating Characteristics-curve (AUROC) and Area Under the Precision Recall-curve (AUPR). We evaluate three CSNI methods by computing the mean AUROC and AUPR across all cells.

The Area Under the Receiver Operating Characteristic-curve (AUROC) and Area Under the Precision-Recall curve (AUPR) metrics are common metrics for evaluating a predicted GRN with a ground-truth GRN [46]. To compare a predicted cell-specific GRN with the ground-truth cell-specific GRN, the top 10’000 interactions per cell is retained, and the mean AUROC and AUPR scores are calculated across all cells.

### Comparison of RNA velocity methods

For each of the 14 different predefined backbones, nine datasets were generated with three difficulty settings and three different seeds. The different difficulty settings were obtained by multiplying the transcription rate by a factor of 25 (easy), 5 (medium) and 1 (hard). After manual quality control, the easy and medium datasets for backbones “bifurcating_converging”, “bifurcating_loop”, “converging” and “disconnected” were removed, resulting in a final collection of 102 datasets.

We calculated two evaluation metrics: the velocity correlation and the velocity arrow cosine. For the velocity correlation, we extracted a ground truth RNA velocity by subtracting for each mRNA molecule the propensity of its production by the propensity of its degradation. If the expression of an mRNA will increase in the future, this value is positive, while it is negative if it is going to decrease. For each gene, we determined its velocity correlation by calculating the Spearman rank correlation between the ground truth velocity with the observed velocity. For the velocity arrow cosine, we determined a set of 100 trajectory waypoints uniformly spread on the trajectory. For each waypoint, we weighted each cell based on a Gaussian kernel on its geodesic distance from the waypoint. These weights were used to calculate a weighted average velocity vector of each waypoint. We then calculated for each waypoint the cosine similarity between this velocity vector and the known direction of the trajectory.

We compared two RNA velocity methods. The velocyto method [22], as implemented in the velocyto.py package, in which we varied the “assumption” parameter between “constant_unspliced” and “constant_velocity”. The scvelo method [23], as implemented in the python scvelo package scvelo.de, in which we varied the “mode” parameter between “deterministic”, “stochastic”, “dynamical”, “dynamical_residuals”. For both methods, we used the same normalized data as provided by dyngen, with no extra cell or feature filtering, but otherwise matched the parameters to their respective tutorial vignettes as well as possible.

To visualize the velocity on an embedding, we used the “velocity_embedding” function, implemented in the scvelo python package.

### Comparison of trajectory alignment with increasing levels of noise

10 base datasets, containing a linear trajectory, were generated using dyngen. For each dataset, cells and experiments were generated twice, resulting in 20 paired datasets containing a similar trajectory with slightly different kinetics. For each of these pairs of datasets, we generated 10 progressively noisier pairs, in which we added noise to the complete count matrix. Let *q* be the 75th quantile of the non-zero values in the count matrix. The noise added to the count matrix is calculated as follows, with *i* being the noise parameter.

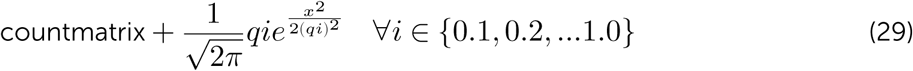

We aligned each of these 100 pairs of trajectories (the first trajectory in the pair with the second trajectory in the pair) using Dynamic Time Warping (DTW) [16]. DTW is designed to align temporal sequences by dilating or contracting the sequences to best match each other. We compared the alignment of DTW with the alignment of DTW after first smoothing the expression along the trajectory using a Gaussian kernel with window size 0.05.

To evaluate a trajectory alignment method on a paired dataset we computed the geodesic distances of each cell from the start of the trajectory. The geodesic distances between the pair of datasets are directly comparable to each other, and thus cells from the first trajectory should align relatively closely to cells from the second trajectory. After aligning the cells, we calculate the mean absolute difference between the geodesic distances of the pairs of cells, which should ideally be 0.

