## Supplementary Figures for "dyngen: a multi-modal simulator for spearheading new single-cell omics analyses"

Table S1: **Feature comparison of single-cell synthetic data generators and simulators.** [1] As showcased by vignette. [2] As showcased by Van den Berge et al. [1].

| | splatter | powsimR | PROSSTT | SymSim | dyngen |
|---|---|---|---|---|---|
| **Available modality outputs** | | | | | |
| - mRNA expression | ✓ | ✓ | ✓ | ✓ | ✓ |
| - Pre-mRNA expression | | | | | ✓ |
| - Protein expression | | | | | ✓ |
| - Promotor activity | | | | ✓ | ✓ |
| - Reaction activity | | | | | ✓ |
| **Available ground-truth outputs** | | | | | |
| - True counts | ✓ | | | ✓ | ✓ |
| - Cluster labels | ✓ | ✓ | | ✓ | ✓ |
| - Trajectory | ✓ | | ✓[1] | ✓ | ✓ |
| - Batch labels | ✓ | | | | ✓[1] |
| - Differential expression | | | | ✓ | ✓[2] |
| - Knocked down regulators | | | | | ✓ |
| - Regulatory network | | | | ✓ | ✓ |
| - Cell-specific regulatory network | | | | | ✓ |
| **Emulate experimental effects** | | | | | |
| - Single-cell RNA sequencing | ✓ | ✓ | ✓ | ✓ | ✓ |
| - Batch effects | ✓ | | | | ✓[1] |
| - Knockdown experiment | | | | | ✓ |
| - Time-series | | | | | ✓ |
| - Snapshot | | | | | ✓ |
| **Evaluation applications** | | | | | |
| - Clustering | ✓ | ✓ | | ✓ | |
| - Trajectory inference | ✓ | | ✓ | ✓ | ✓ |
| - Network inference | | | | ✓ | ✓ |
| - Cell-specific network inference | | | | | ✓ |
| - Differential expression | ✓ | | | | ✓[2] |
| - Batch effect correction | ✓ | | | | ✓ |
| - RNA Velocity | | | | | ✓ |
| - Trajectory alignment | | | | | ✓ |

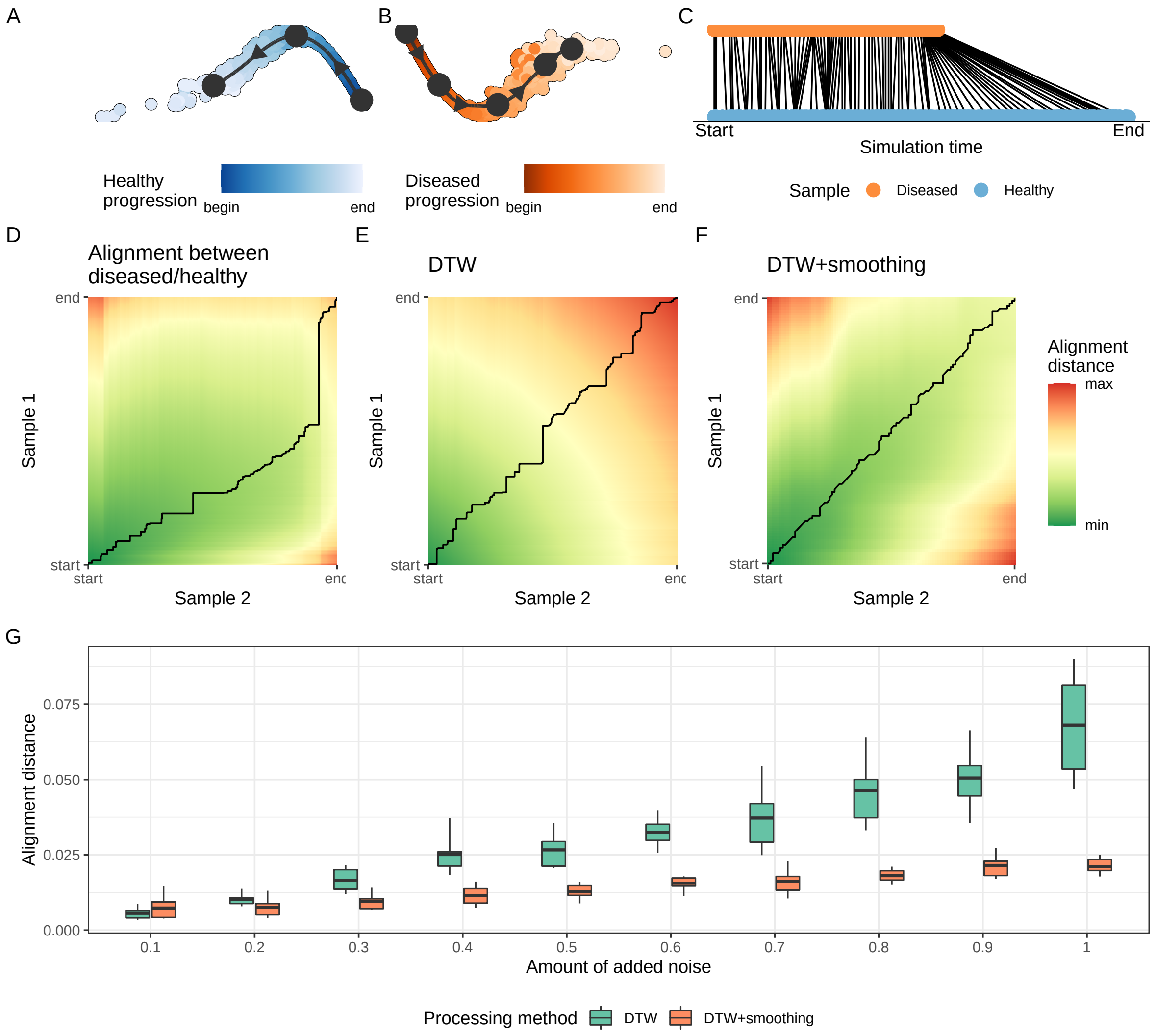


Figure S1: **dyngen allows benchmarking of trajectory alignment methods. A, B**: Two samples containing a linear trajectory, generated by dyngen. **C, D**: Result of the DTW alignment on the samples in **A** and **B**. In **C**, the individual mappings of the alignment between cells are shown. In **D**, the accumulated distance matrix between the two trajectories is shown, including the black warping path, corresponding to these cell to cell alignments in **C**. **E, F**: Shows the accumulated distance matrices obtained after using DTW on two trajectories where noise (noise level of 0.4) was added to the count matrix. In E the complete count matrices were used to perform the alignment. In F, smoothed pseudocells were used. **G:** Shows the influence of added noise to the different processing methods. We can see that DTW + smoothing performs best in noisy circumstances.

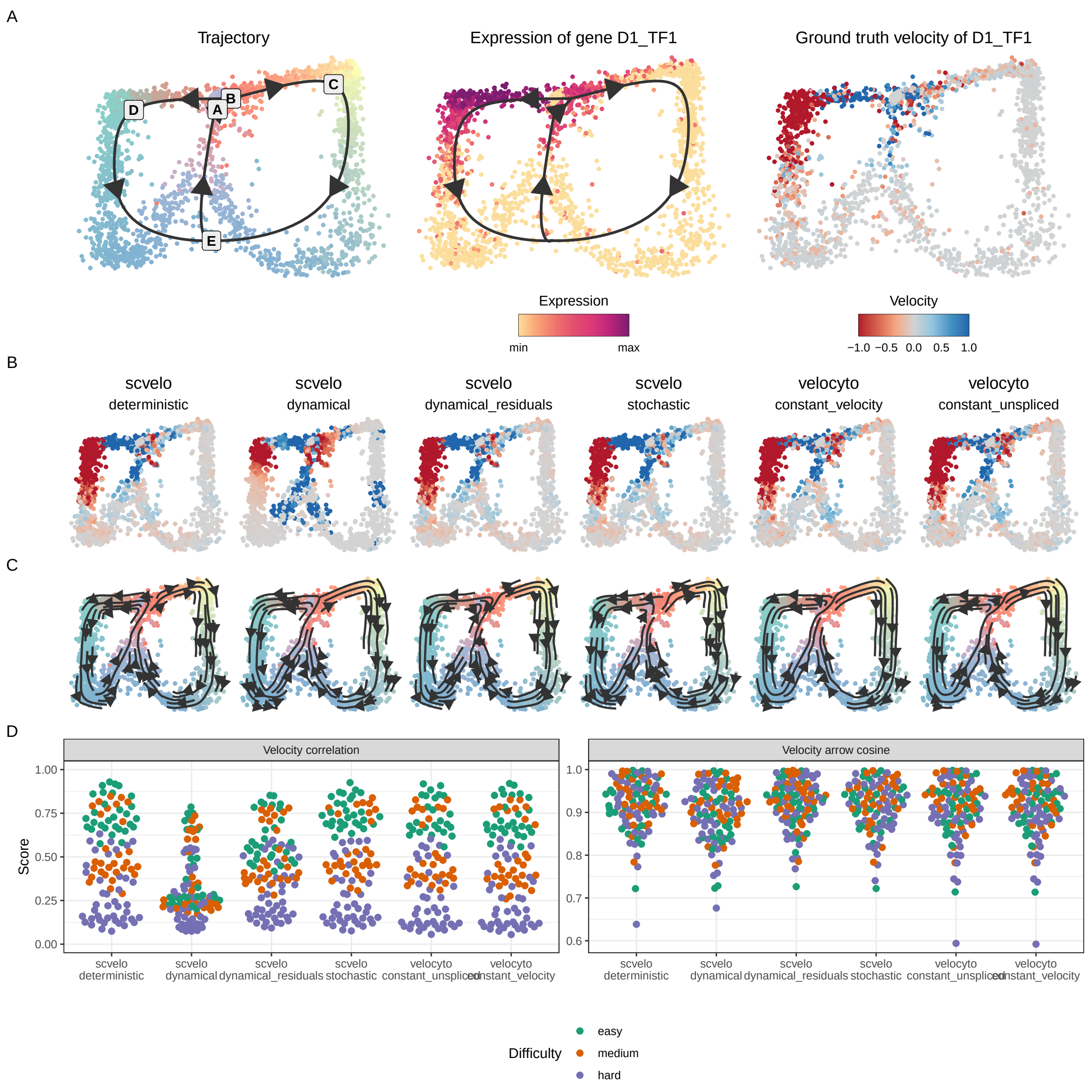


Figure S2: **dyngen allows benchmarking of RNA velocity methods. A:** An example bifurcating cycle dataset, with as illustration the expression and ground truth velocity of a gene D1_TF1 that goes up and down in one branch of the trajectory. **B:** The RNA velocity estimates of gene D1_TF1 by the different methods. **C:** The velocity stream plots produced from the predictions of each method, as generated by scvelo. **D:** The predictions scored by two different metrics, the velocity correlation and the velocity arrow cosine. The velocity correlation is the correlation between the ground-truth velocity (A, right) and the predicted velocity (B). The velocity arrow cosine is the cosine similarity between the direction of segments of the ground-truth trajectory (A, left) and the RNA velocity values calculated at those points (C).

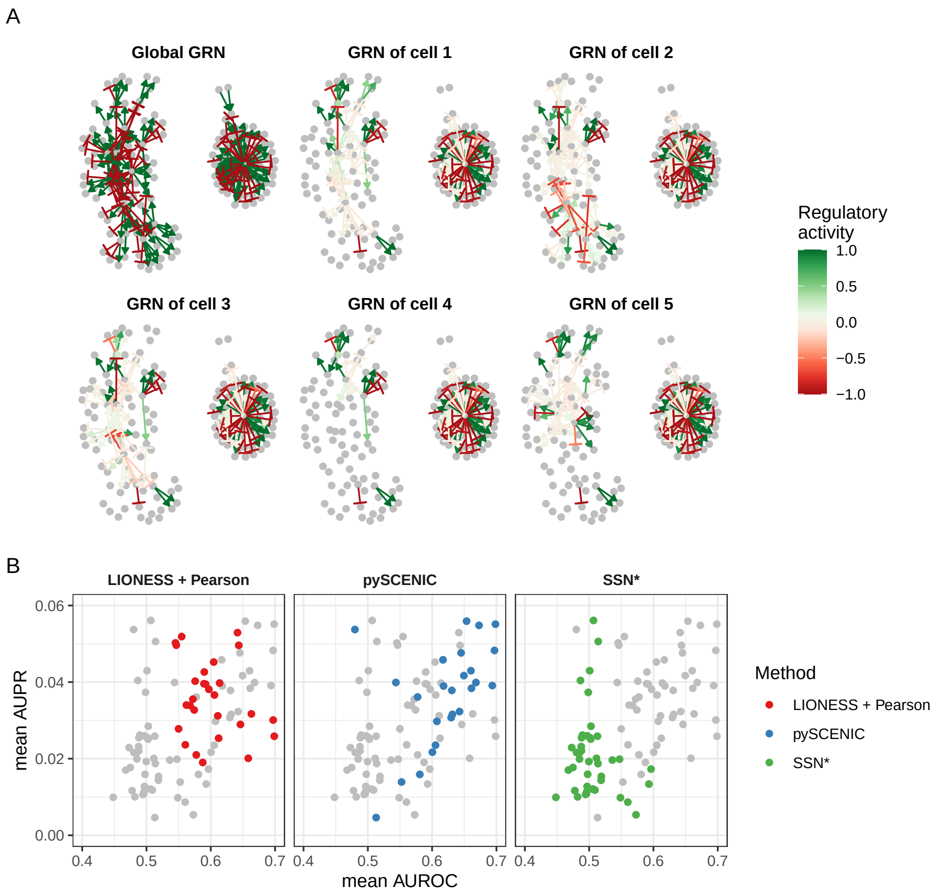


Figure S3: **dyngen allows benchmarking Cell-specific Network Inference (CSNI) methods. A:** A cell is simulated using the global gene regulatory network (GRN, top left). However, at any particular state in the simulation, only a fraction of the gene regulatory interactions are active. **B:** CSNI methods were executed to predict the regulatory interactions that are active in each cell specifically. Using the ground-truth cell-specific GRN, the performance of each method was quantified on 14 dyngen datasets.

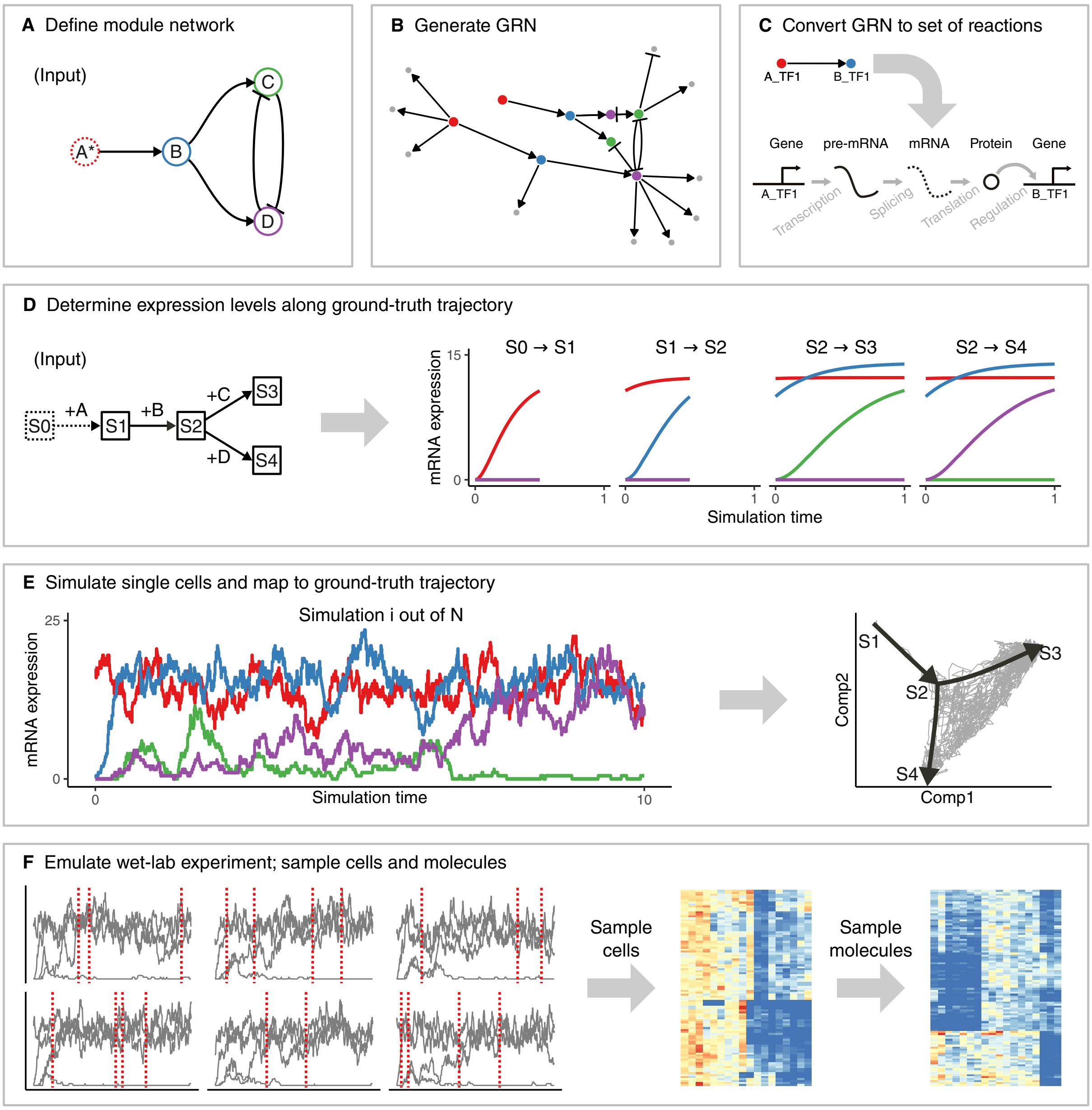


Figure S4: **The workflow of dyngen consists of six main steps. A:** The user needs to specify the desired module network or use a predefined module network. The module network is what determines the dynamic behaviour of simulated cells. **B:** The number of desired transcription factors (which drive the desired dynamic process) are amongst the given modules and adds regulatory interactions according to the module network. Additional target genes (which do not influence the dynamic process) are added by sampling interactions from GRN interaction databases. **C:** Each gene regulatory interaction in the GRN is converted to a set of biochemical reactions. **D:** Along with the module network, the user also needs to specify the backbone structure of expected cell states. The average expression of each edge in the backbone is simulated by activating a restricted set of genes for each edge. **E:** Multiple Gillespie SSA simulations are run using the reactions defined in step C. The counts of each of the molecules at each time step are extracted. Each time step is mapped to a point in the backbone. **F:** The molecule levels of multiple simulations are shown over time (left). From each simulation, multiple cells are sampled (from left to middle). Technical noise from profiling is simulated by sampling molecules from the set of molecules inside each cell (from middle to right).

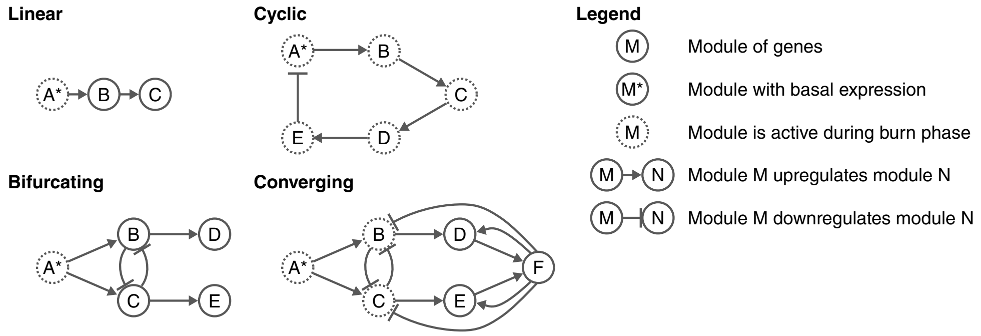


Figure S5: **The module network determines the type of dynamic process which simulated cells will undergo.** A module network describes the regulatory interactions between sets of transcription factors which drive the desired dynamic process.

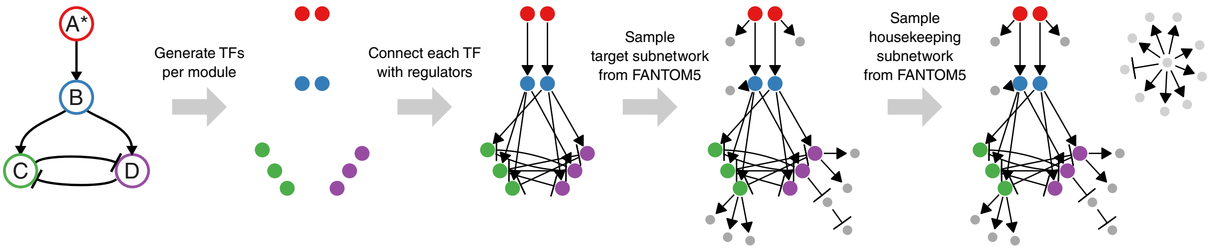


Figure S6: **Generating the feature network from a backbone consists of four main steps.**

Table S2: **Reactions affecting the abundance levels of pre-mRNA $x_G$, mature mRNA $y_G$ and proteins $z_G$ of gene $G$.** Define the set of regulators of $G$ as $R_G$, the set of upregulating regulators of $G$ as $R_G^+$, and the set of down-regulating regulators of $G$ as $R_G^-$. Parameters used in the propensity formulae are defined in Table S3.

| Reaction | Effect | Propensity |
|---|---|---|
| Transcription | $\emptyset \rightarrow x_G$ | $\text{xpr}_G \times \frac{\text{bas}_G - \text{ind}_G^{\lvert R_G^+ \rvert} + \prod_{H \in R_G^+} (\text{ind}_G + \text{bind}_{G,H})}{\prod_{H \in R_G} (1 + \text{bind}_{G,H})}$ |
| Splicing | $x_G \rightarrow y_G$ | $\text{ysr}_G \times x_G$ |
| Translation | $y_G \rightarrow y_G + z_G$ | $\text{zpr}_G \times y_G$ |
| Pre-mRNA degradation | $x_G \rightarrow \emptyset$ | $\text{ydr}_G \times x_G$ |
| Mature mRNA degradation | $y_G \rightarrow \emptyset$ | $\text{ydr}_G \times y_G$ |
| Protein degradation | $z_G \rightarrow \emptyset$ | $\text{zdr}_G \times z_G$ |

Table S3: **Default parameters defined for the calculation of reaction propensity functions.**

| Parameter | Symbol | Definition |
|---|---|---|
| Transcription rate | $\text{xpr}_G$ | $\in U(10, 20)$ |
| Splicing rate | $\text{ysr}_G$ | $= \ln(2)\ /\ 2$ |
| Translation rate | $\text{zpr}_G$ | $\in U(100, 150)$ |
| (Pre-)mRNA half-life | $\text{yhl}_G$ | $\in U(2.5, 5)$ |
| Protein half-life | $\text{zhl}_G$ | $\in U(5, 10)$ |
| Interaction strength | $\text{str}_{G,H}$ | $\in 10^{U(0,2)}$ * |
| Hill coefficient | $\text{hill}_{G,H}$ | $\in U(0.5, 2)$ * |
| Independence factor | $\text{ind}_G$ | $\in U(0, 1)$ * |
| (Pre-)mRNA degradation rate | $\text{ydr}_G$ | $= \ln(2)\ /\ \text{yhl}_G$ |
| Protein degradation rate | $\text{zdr}_G$ | $= \ln(2)\ /\ \text{zhl}_G$ |
| Dissociation constant | $\text{dis}_H$ | $= 0.5 \times \frac{\text{xpr}_H \times \text{ysr}_H \times \text{zpr}_H}{(\text{ydr}_H + \text{ysr}_H) \times \text{ydr}_H \times \text{zdr}_H}$ |
| Binding strength | $\text{bind}_{G,H}$ | $= \text{str}_{G,H} \times (\text{z}_H\ /\ \text{dis}_H)^{\text{hill}_{G,H}}$ |
| Basal expression | $\text{bas}_G$ | $= \begin{cases} 1 & \text{if } \text{R}_G^+ = \emptyset \\ 0.0001 & \text{if } \text{R}_G^- = \emptyset \text{ and } \text{R}_G^+ \neq \emptyset\ * \\ 0.5 & \text{otherwise} \end{cases}$ |

*: unless $G$ is a TF, then the value is determined by the backbone.

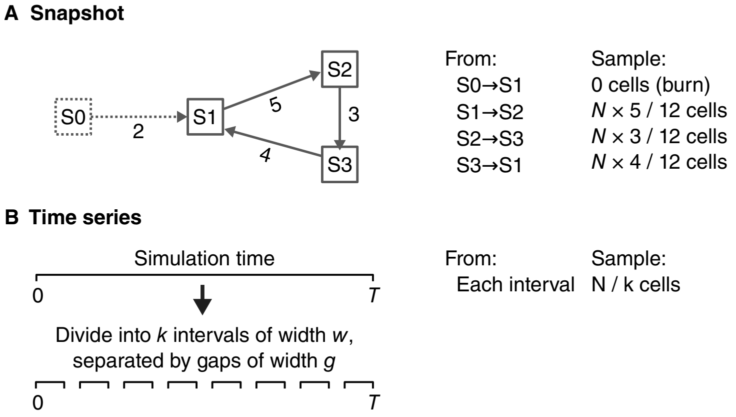


Figure S7: **Two approaches can be used to sample cells from simulations: snapshot and time-series.**

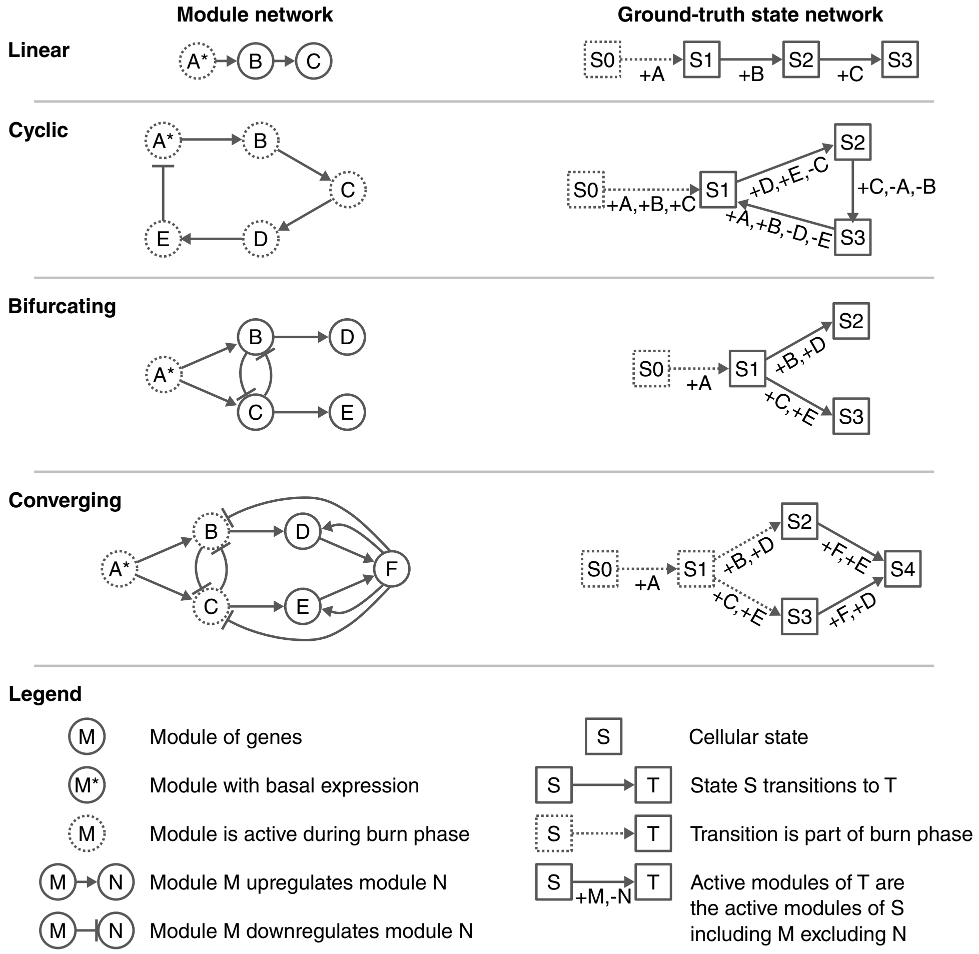


Figure S8: **Examples of the ground-truth state networks which need to be provided alongside the module network.**
